# Mechanics Of Flight Feathers: Effects Of Captivity?

**DOI:** 10.1101/2020.04.14.038877

**Authors:** Steven J. Portugal, Robert L. Nudds, Jonathan A. Green, R. McNeil Alexander, Patrick J. Butler, Robert F. Ker

## Abstract

Feathers act as aerodynamic cantilevers, and to withstand the prolonged cyclical loading that occurs during flight, feathers must be stiff, lightweight and strong. We experimentally tested the differences in feather structure, primarily stiffness and size, between (*a*) wild and captive Barnacle Geese *Branta leucopsis,* and (*b*) primary feathers dropped during the annual flight feather moult, and those feathers freshly regrown during the moult process. We found that, despite having undergone a 5,000km round-trip migration, flight feathers dropped during moult in the wild geese were stiffer than those measured in the captive geese, both for those dropped during moult and those re-grown. We propose that this may be related to diet or stress in the captive geese.

## INTRODUCTION

Birds use feathers for insulation, display, and camouflage. Most importantly, is the crucial role they play in flight. Feathers act as aerodynamic cantilevers, and to withstand the prolonged cyclical loading that occurs during flight, feathers must be stiff, lightweight and strong (Bonser 1996; Weber *et al*. 2005). This cyclic loading can be extreme. For a small passerine, for example, this can involve over 40 million wing-beats during the completion of one migratory journey from sub-Saharan Africa to western-Europe (Weber *et al*. 2005). The loss or breakage of flight feathers has significant deleterious effects on such flights, and on overall flight performance in general (Verbeek and Morgan 1980; Tucker 1991). Therefore, it is vital for birds to synthesise good quality feathers and in particular, for them to have good bending stiffness to avoid breakage. Although the actual mechanical properties of the feathers have only rarely been studied (Bonser and Purslow 1995; Bonser 1996; Corning and Biewener 1998; Weber *et al*. 2005), it is established that bending stiffness is a vital aspect in making the feather act as an effective aerofoil (Bonser 1996; Weber *et al*. 2005).

If breakage or loss of feathers should occur, birds typically have to wait until the next flight feather moult before the feather can be replaced, as unlike analogous tissues such as bones and human fingernails, feathers do not have the capacity for self-repair (e.g. Weber *et al*. 2005). The quality of the feather produced during flight feather moult is vital to many aspects of the life-history of birds, with poorer quality feathers having been shown to result in reduced over-winter survival rates and subsequent reduced breeding success the following year for surviving individuals (Dawson *et al*. 2000). However, while we know that moult and feather replacement are an extremely significant aspect of the annual cycle, as highlighted by Weber *et al*. (2005), there are very few studies which experimentally demonstrate the mechanical advantages of new flight feathers compared to visibly undamaged but old plumage (see Chai and Dudley 1999; Williams and Swaddle 2003). Here, by measuring bending stiffness and thickness of captive Barnacle Geese *(Branta leucopsis)* feathers, we present opportunistically-collected data that provides an estimate of the structural benefit gained from the re-growth of flight feathers during moult. We predicted that new feathers grown during the moult process would be stiffer than those feathers dropped at the onset of moult; the reduction of stiffness in the moulted feathers being a result of wear and tear. We also provide an estimate of the impact that migratory flights have on feather bending stiffness, by comparing flight feathers dropped during moult in these adult captive birds that had never flown with those from wild migratory barnacle geese that had completed a 5000 km round trip from Svalbard to Caerlaverock, south-west Scotland, and back (see Butler *et al*. 1998). We predicted that the flight feathers dropped during moult in the captive birds would be stiffer than those dropped by the wild moulting Barnacle Geese, as they have not been used during sustained periods of flight.

## METHODS

### Feather samples

A captive population of Barnacle Geese was maintained at the University of Birmingham, UK. All husbandry details can be found in Portugal *et al*. (2007; 2011a; 2011b). Flight feathers without visible fault bars were collected daily from the aviary during the wing moult period of the captive geese in 2008. The geese were observed regularly to assess moult stage and only the 9^th^ primary was used for analysis, identified by the total length of the feather. On the completion of moult, the newly grown 9^th^ primary feather was removed. Flight feathers from wild Barnacle Geese were collected during wing moult, at Ny-Ålesund, Spitsbergen (78°55’N 11°56’78.917°N 11.933°E,)(see Portugal *et al*. 2011c; Portugal *et al*., 2019a; Portugal *et al,.* 2019b). It was possible to check the area twice daily, ensuring feathers were freshly dropped and had not been lying on the ground for any extended time period. After collection, feathers of both the captive and wild geese were weighed immediately, and the length of the feather recorded. The diameter of the calamus and the dorso-ventral thickness were also measured before the samples were frozen at −20°C.

### Mechanical testing equipment

The elastic constant (*k* in Nmm^-1^) of the feathers was measured in a three-point bending set-up, using an Instron 8500 dynamic testing machine. The feather rested on two aluminium supports on the ventral side, which were attached to the actuator via a steel plate. The third contact point, on the dorsal surface was stationary and fixed to the load cell. The spacing of the supports was 20 cm. Each feather was clamped into place between two cable-clamps screwed firmly together to avoid any movement or twisting during the experimental procedure. For measurements of bending stiffness on the calamus, the feather was clamped at one end by the very tip of the calamus, and then again at the calamus-rachis edge. For measurements of bending stiffness from the rachis, the feather was clamped again at the calamus-rachis edge, and then three quarters down the length of the rachis. The point of bending was 15 cm along the length of the rachis, and 1 cm on the calamus.

Tests were performed in air at room temperature (24.2 ± 2.1° C). A pre-load of – 12 N was applied to the calamus, and – 3 N for the rachis. Haversine oscillations (to determine the distance between two points on a sphere) were applied at a frequency of 5 Hz under position control, with an amplitude of 0.5 mm for the calamus, and 1 mm for the rachis. The load cell output was set to zero when the feather specimen was not in contact with the dorsal support.

### Data Analyses

When a feather was under load, the relationship between force (*F*, N) and extension (*e*, mm) was curvi-linear and the feathers behaved viscoelastically: the bending tension curve differed from the unloading curve producing a hysteresis loop (Fig. 1). Therefore, the feathers undergoing bending do not conform to Hooke’s law where a structure’s length should alter by an amount proportional to the force applied and hence in the relationship:

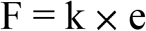

*k* should be constant. To approximate *k* a simple least squares regression (*r*^2^ > 0.999 in all cases) was fitted through the resulting hysteresis curves (a single line through the loading and unloading curve). This was carried out for each test sample. As the samples were pre-loaded the intercept term is uninformative. The slopes (*k*) of the resulting regression equations alone were tested for a difference between feather part (rachis or calamus) and the three feather cohorts: regrown feathers from captive birds (captive re-grown feathers) (*N* = 9), dropped feathers from captive birds (captive dropped feathers) (*N* = 16) and dropped feathers from wild birds (wild dropped feathers) (*N*= 17) using a GLM with sample dorso-ventral thickness (mm) included as a covariate. An interaction term (feather part × cohort) was also included in the ANOVA to determine whether the pattern of *k* across feather parts was consistent among cohorts. Non-significant terms were stepwise deleted from the GLM and the GLM rerun until all remaining independent variables were significantly affecting *k*. A two-way ANOVA was used to determine whether feather thickness differed between cohort and feather part. Bonferroni post-hoc tests were used to locate differences between the cohorts and partial (_p_) eta squared (*η*_p_^2^) is used as an estimate of effect size throughout the analyses. Statistical analyses were performed using IBM^®^ SPSS^®^ Statistics v.20.

**Figure 1.**
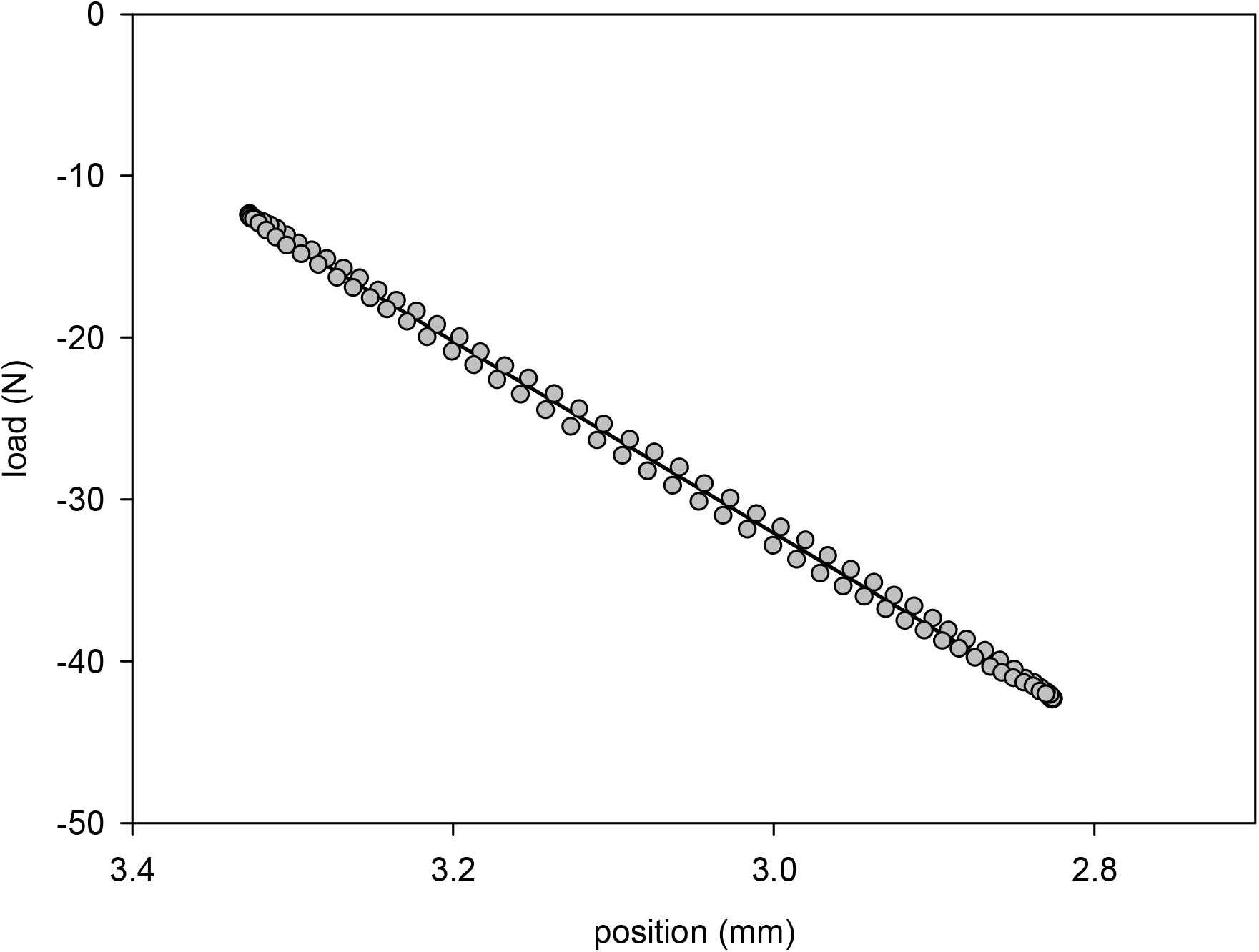
A sample of a typical hysteresis loop during a loading cycle of a primary feather. These data are from the calamus section of a newly re-grown primary feather, taken from a captive Barnacle goose. Linear regression, when applied to the loop, represents a good approximation to the major axis of the fatigue ellipse.

## RESULTS

There was no interaction between cohort and feather part (*F*_2, 77_ = 1.144, *η*_p_^2^ = 0.03, *P* = 0.324) and sample thickness had no effect on k, the feather stiffness constant (*F*_1, 79_ = 0.172, *η*_p_^2^ < 0.01, *P* = 0.679). The final GLM, with the interaction term and covariate thickness removed, showed *k* to differ between both cohort and feather part (cohort: *F*_2, 80_ = 8.1, *η*_p_^2^ = 0.17, *P* = 0.001; feather part: *F*_1, 80_ = 496.1, *η*_p_^2^ = 0.86, *P* < 0.001; Table 1). The Bonferroni post-hoc test showed that the difference between cohorts was due to the wild dropped cohort having higher values of *k* (a greater stiffness) than the two captive cohorts. The difference (*P* = 0.003) in *k* between captive dropped and wild dropped was −5.18 ± 3.76 N mm^-1^ (mean difference ± 95 C.I.), and the difference (*P* = 0.003) between captive regrown, and wild dropped −6.17 ± 4.45 N mm^-1^ (mean difference ± 95 C.I.) (Table 1). *k* was similar (P = 1.000) for captive dropped and captive regrown feather samples, 1.00 ± 3.66 N mm^-1^ (mean difference ± 95 C.I.). The standardised residuals generated by the three GLMs were normally distributed in each case (Kolmogorov-Smirnov test: *P* > 0.200).

**Table 1.** Mean (± S.E.M.) slopes (*k* in N mm^-1^) extracted from the hysteresis curves during stiffness testing from the rachis and calamus sections of the primary feathers of captive and wild Barnacle Geese.

| Feather part | Captive re-grown | Captive dropped | Wild dropped |
| --- | --- | --- | --- |
| Rachis | $15.94 \pm 0.83$ | $17.83 \pm 0.96$ | $21.14 \pm 0.80$ |
| Calamus | $46.17 \pm 2.94$ | $46.25 \pm 2.11$ | $53.30 \pm 1.84$ |

Although the calamus was consistently stiffer than the rachis, thickness had no effect on *k* within each cohort and feather part, which is presumably due to the narrow range of thicknesses within each group. Nonetheless, it was a surprising result, which warranted further investigation. The feather calamus was thicker than the rachis; (*F*_1, 78_ = 180.017, *η*_p_^2^ = 0.93, *P* < 0.001) and differed between cohorts (*F*_2, 78_ = 4.626, *η*_p_^2^ = 0.11, *P* = 0.013), but the pattern of difference between the feather parts varied between cohort (*F*_2, 78_ = 1.558, *η*_p_^2^ = 0.20, *P* < 0.001). A Bonferroni post-hoc test indicated that the difference detected in thicknesses between cohorts was due to the wild dropped feathers having thicker feather shafts (*P* = 0.021) than the captive dropped feathers. There was no difference (*P* = 1.000) in feather thickness between the wild dropped and captive regrown cohorts, nor a difference (*P* = 0.066) between captive dropped and captive regrown cohorts. The interaction effect was due to the captive dropped cohort. The drop in feather thickness between the calamus and rachis was very similar in the wild dropped and captive regrown cohorts. The wild dropped cohort, however, had a higher mean calamus thickness and lower mean rachis thickness than the two captive cohorts (Tables 1 and 2). Again, the standardised residuals generated by the two-way ANOVA were normally distributed (Kolmogorov-Smirnov test: *P* > 0.200).

**Table 2.** Mean (± S.E.M.) feather thicknesses in the dorso-ventral plane (mm) of rachis and calamus sections of the primary feathers of captive and wild Barnacle Geese.

| Feather part | Captive re-grown | Captive dropped | Wild dropped |
| --- | --- | --- | --- |
| Rachis | $3.56 \pm 0.12$ | $2.87 \pm 0.11$ | $3.54 \pm 0.08$ |
| Calamus | $6.32 \pm 0.13$ | $6.45 \pm 0.57$ | $6.34 \pm 0.07$ |

To summarise, the wild dropped feathers had both a stiffer rachis and calamus than those of both captive cohorts, which showed similar stiffness. The relative drop in stiffness from the calamus to the rachis was also consistent between the three cohorts. Feather stiffness did not relate to feather thickness, with the wild dropped cohort having the stiffest, yet not thickest calamus (Tables 1 and 2). The thickness of the calamus relative to the rachis was greater for the captive dropped cohort than for the captive regrown and wild dropped cohorts, which were similar.

## DISCUSSION

Contrary to our predictions, the dropped flight feathers of the wild geese were significantly stiffer than the feathers in the captive birds. These stiffer feathers in the wild geese are despite the birds having undergone a 5000 km round trip from Svalbard to Caerlaverock (Butler *et al*. 1998) between flight-feather moults, in comparison to the captive geese which had never flown. This suggests that there are fundamental differences in the structure of the feathers grown by captive geese, compared to those produced by wild birds. This is further supported by the previously demonstrations of the negative impact that migration and flight can have on feather fatigue and stiffness, and the subsequent influence this can have on moult strategies (Weber *et al*. 2005). Thus, it would be expected that the flight feathers dropped by the wild geese should have a reduced feather stiffness compared to captive birds. Reduced feather stiffness can reduce flight performance for many reasons, including diminishing the aerodynamic force created during flapping flight (Nordberg, 1985), and has been demonstrated to impair escape-flight performance in European Starlings *(Sturnus vulgaris*). It is likely, therefore, that if it had been possible to collect the freshly re-grown flight feathers from the wild Barnacle Geese, these feathers would have been even stiffer than those which were dropped during moult, exacerbating the differences between the captive and wild birds.

Captivity can have pronounced effects on external, skeletal and soft-tissue morphological traits (Courtney-Jones *et al*. 2018, and references therein). The observed disparity in stiffness between the feathers of wild and captive birds may, therefore, potentially be explained by two primary factors related to captivity, (*i*) diet, and (*ii*) stress. Nitrogen requirements are elevated during moult, due to the synthesis of new feathers (Murphy and King, 1992) and many wild species of waterfowl preferentially switch their diet during flight-feather moult and select the nitrogen-rich tips of certain vegetation types (Fox and Kahlert, 1999). It is possible, that even though food was available *ad libitum* to the captive geese in the present study that it did not provide sufficient nitrogen for efficient feather synthesis. Stress can cause abnormalities in feather development during moult (DesRochers *et al*. 2009). The mechanism behind this phenomenon is the interaction between stress, the release of corticosterone (CORT), and protein inhibition (Romero *et al*., 2005). CORT is released into the blood stream by avian adrenal tissue, typically in response to a stressor (Holmes and Phillips, 1976; Romero *et al*. 2005). CORT has degradative effects on proteins and acts to suppress protein synthesis. In most birds, predictable (i.e. not following a stochastic event) CORT concentrations are lower during moult than at any other time of the year (Romero 2002; DesRochers *et al*. 2009). As feathers are comprised primarily of the protein keratin, it is likely this suppression in CORT during moult is to avoid the protein catabolic activity of CORT from inhibiting the protein deposition necessary to produce feathers (Sapolsky *et al*. 2000; Romero 2002). DesRochers *et al*. (2009), for example, demonstrated that birds which had higher circulating levels of CORT had lighter feathers, reduced feather strength, and even a different feather microstructure. It is possible, therefore, that the lower feather stiffness values for the captive birds, in comparison to the wild geese, may be linked to higher stress levels, and as a result, a lack of suppression of CORT during moult. The source of stress in the captive geese, and the associated potential increase in circulating levels of CORT, could be linked to either (*i*) disturbance or (*ii*) artificial light. During moult, the captive geese were involved in a series of behavioural and physiological experiments (see Portugal *et al*. 2007, 2009a, 2009b), which may have elevated CORT levels. Similarly, CORT levels may have been elevated due to the urban environment the geese were living in, and associated artificial light. Melatonin is one of the fundamental hormones involved in the regulation of daily physiological cycles and is released at night and supressed by daylight (Gwinner, 1996). Natural rhythms in melatonin release can, however, be interrupted by artificial light (Le Tallec *et al*. 2013). Moreover, it has been demonstrated that artificial interruptions to the typical circadian rhythm in melatonin can act as a stressor, and thus increase CORT levels (Xu *et al*. 2016). It is possible, consequently, that a combination of handling, experiments, and artificial light interacting with circulating CORT and melatonin may be the underlying cause for the poorer feather stiffness in the captive geese compared to their wild counterparts.

While potential stress and diet may account for the differences in feather stiffness between the wild and captive birds, it does not explain the similarities in feather stiffness values between the captive dropped and captive regrown. Although the birds had never flown, thus not experienced the feather-fatigue associated with prolonged power flight (Weber *et al*., 2006), the feathers of the geese would still have been used for thermoregulation, exposed to UV-radiation, and been subjected to attack from parasites. Thus, it would have been expected that there would be some evidence of degradation over time, and consequently a decrease in stiffness in the moulted feathers. The lack of difference may be a result of a combination of a lower feather parasite load in captivity in comparison to wild conspecifics, and feather degradation from parasites affecting feather parameters other than stiffness. Interestingly, heart rates measured during flight in captive geese are typically higher than those recorded in wild birds (Butler *et al*., 2000), and a potential link between reduced feather stiffness in captive birds and increased flight costs is worthy of further investigation.

We are grateful to Richard Bonser for useful discussions and to Maarten Loonen for assistance with feather collection. Funding for this study was provided by a NERC grant to P.J.B, while S.J.P was funded by a BBSRC studentship. Conceptualisation, S.J.P., P.J.B. and R.F.K.; methodology, S.J.P., R.M.A. and R.F.K.; resources, R.M.A. and R.F.K.; data collection, S.J.P. and R.F.K., formal analysis, R.L.N. and J.A.G.; writing, reviewing and editing, S.J.P., R.L.N., J.A.G., P.J.B. and R.F.K.

